# Identification of proteins that bind extracellular microRNAs secreted by the parasitic nematode *Trichinella spiralis*

**DOI:** 10.1101/2023.05.29.542739

**Authors:** A. Brown, M.E. Selkirk, P. Sarkies

## Abstract

Small non-coding RNAs such as microRNAs (miRNAs) are conserved across eukaryotes and play key roles in regulating gene expression. In many organisms, miRNAs are also secreted from cells, often encased within vesicles such as exosomes and sometimes extravesicular. The mechanisms of miRNA secretion, how they are stabilised outside of cells and their functional importance are poorly understood. Recently we characterised the parasitic nematode *Trichinella spiralis* as a model to study miRNA secretion. *T. spiralis* larvae secrete abundant miRNAs which are largely extravesicular. Here, we investigated how *T. spiralis* miRNAs might remain stable outside of cells. Using proteomics, we identified two RNA binding proteins secreted by *T. spiralis* larvae and characterised their RNA binding properties. One, a homologue of the known RNA binding protein KSRP, binds miRNA in a selective and sequence-specific fashion. Another protein, which is likely a novel RNA binding protein, binds non-selectively to miRNA. Our results suggest a possible mechanism for miRNA secretion by *T. spiralis* and may have relevance for understanding the biology of extracellular miRNA more widely.

## Introduction

Small (16-36nt) non-coding RNAs (sRNAs) are key regulators of gene expression conserved across eukaryotes. Different pathways generate functionally distinct classes of sRNAs[1] but in general sRNAs associate with Argonaute proteins, which catalyse efficient recognition of target sites within RNAs through sense-antisense base pairing[2]. This then usually results in downregulation of the target RNA. microRNAs are one of the most abundant classes of sRNAs[3]. miRNAs are conserved in all animals and their activity has been demonstrated to be essential for successful development in several organisms[4]. miRNAs also play important roles in maintaining gene expression in differentiated cells and transcriptional responses to external stimuli. Target recognition by miRNA/Argonaute complexes leads to gene expression changes by inducing mRNA degradation and through disrupting translation[5].

Intracellular functions and mechanisms of miRNAs have been extensively characterised. However, much more mysterious is whether miRNAs might have functions outside cells. Interest in this area began with the unambiguous demonstration that sRNAs, including miRNAs, are transported between tissues in plants[6] and nematodes[7]. In animals, secretion of extracellular miRNAs has been observed from a wide variety of different tissues and cultured cells, and stable extracellular miRNAs have been identified in many different extracellular fluids such as saliva, breast milk and urine[8]. There has been much speculation that these extracellular RNAs could be important in cell-to-cell communication. However, evidence for cell-to-cell transfer of miRNAs in animals is limited and there are many doubts over whether extracellular miRNAs can be delivered in sufficient quantities to exert changes in gene expression[9]. Importantly, there is also limited evidence that extracellular miRNAs are bound to Argonaute proteins, thus exactly how they would integrate into gene expression control mechanisms in recipient cells is unclear.

Lack of understanding about the functions of extracellular miRNAs is accompanied by considerable uncertainty over the mechanism whereby intracellular miRNAs are targeted for secretion and stabilised when in the extracellular environment[9]. The dominant theory has been that miRNAs are enclosed within exosomes which protects them from extracellular nuclease activity[10]. Some intracellular sorting proteins have been implicated in selecting miRNAs for export via this pathway[10]. Although this is the most straightforward way to explain how stable miRNAs can exist outside of cells in the absence of Argonaute proteins, it is notable that up to 50% of mammalian extracellular miRNAs are not enclosed in vesicles[11,12], so there may be other mechanisms involved. How these miRNAs remain stable is poorly understood[9].

Parasitic nematodes have emerged as an interesting model to study the mechanism and function of extracellular RNAs[13,14]. Several species of parasitic nematodes secrete sRNAs, including abundant miRNAs. Similarly to mammals, a substantial fraction of secreted miRNAs are enclosed within vesicles[15]. Some evidence exists that miRNAs secreted by parasitic nematodes in vesicles could be taken up by host cells and potentially contribute to gene regulation[16]. Interestingly, an Argonaute protein has been shown to be secreted by the parasitic nematode *Heligmosomoides polygyrus*[17]. However, this Argonaute protein binds a different class of small non-coding RNAs known as 22G-RNAs[17] so is unlikely to be involved in stabilisation or delivery of miRNAs.

Recently we developed the parasitic nematode *Trichinella spiralis* as a model system to study extracellular small non-coding RNAs. *T. spiralis* is unusual because its life cycle comprises both intracellular and extracellular parasitic phases. Adults mate and produce offspring as extracellular parasites in the gut, but the larval offspring migrate to the muscle cells of the host where they encyst as an intracellular parasite. The muscle stage larvae remain in this state until the animal is predated on, whereby they are released in the digestive tract and develop into adults to complete the life cycle[18]. Infection by *T. spiralis* larvae leads to a number of changes in muscle cells, most notably cell cycle re-entry and extensive remodelling[19]. These processes may involve direct manipulation of gene expression by factors secreted by *T. spiralis* larvae[20], which include abundant small non-coding RNAs [21]. Interestingly, *T. spiralis* muscle stage larvae (MSL) secrete miRNAs that are almost exclusively not contained within vesicles, whilst adult *T. spiralis* secrete predominantly vesicular miRNAs[21].

In this work we investigate the mechanism of secretion of extravesicular miRNAs by *T. spiralis*. In particular we focus on the question of how secreted miRNAs are stabilised. Using proteomics we discover two secreted RNA binding proteins, one of which is from a protein family never previously implicated in nucleic acid interactions. We show that these proteins bind miRNAs both in vitro and in *T. spiralis* larval secretomes. One protein binds non-selectively to miRNAs and the other binds only to a subset of miRNA. Together our work provides new insights into how extracellular miRNAs are stabilised in parasitic nematodes and may have implications for understanding the mechanisms of miRNA secretion in these organisms.

## Results

### T. spiralis secretome contains RNA binding proteins

We previously showed that *T. spiralis* larvae secrete abundant miRNAs that are not enclosed in vesicles, leading to the question of how these miRNAs might be protected from nuclease activity[21]. We speculated that RNA binding proteins might be secreted alongside miRNAs and that these proteins might bind and stabilise miRNAs. We performed proteomics from secreted material from both adult and muscle stage larval (MSL) *T. spiralis* (Supplemental table 1). We identified subsets of proteins that were enriched in secreted material relative to whole worm extracts (Fig 1A). Although there was a significant overlap between proteins enriched in adult and larval stage secretomes, some proteins were specifically enriched in the MSL secretome (Fig 1B and Fig 1C), indicating that they may be involved in stabilising extracellular miRNAs. We searched all proteins that were present in the secreted material from a manually curated list of RNA binding domains (Supplemental table 2). Several candidate proteins were identified which were enriched in MSL secreted material compared to adult secreted material (Fig 1D; Supplemental table 3). We selected two of these proteins for further characterisation. One, which we refer to as TsPUF, had a region with weak similarity to the Pumilio homology (Puf) domain (Fig 1E) [22]. The other, which we refer to as TsKSRP, contained several matches to the KH domain present in many RNA binding proteins [23].

**Figure 1.**
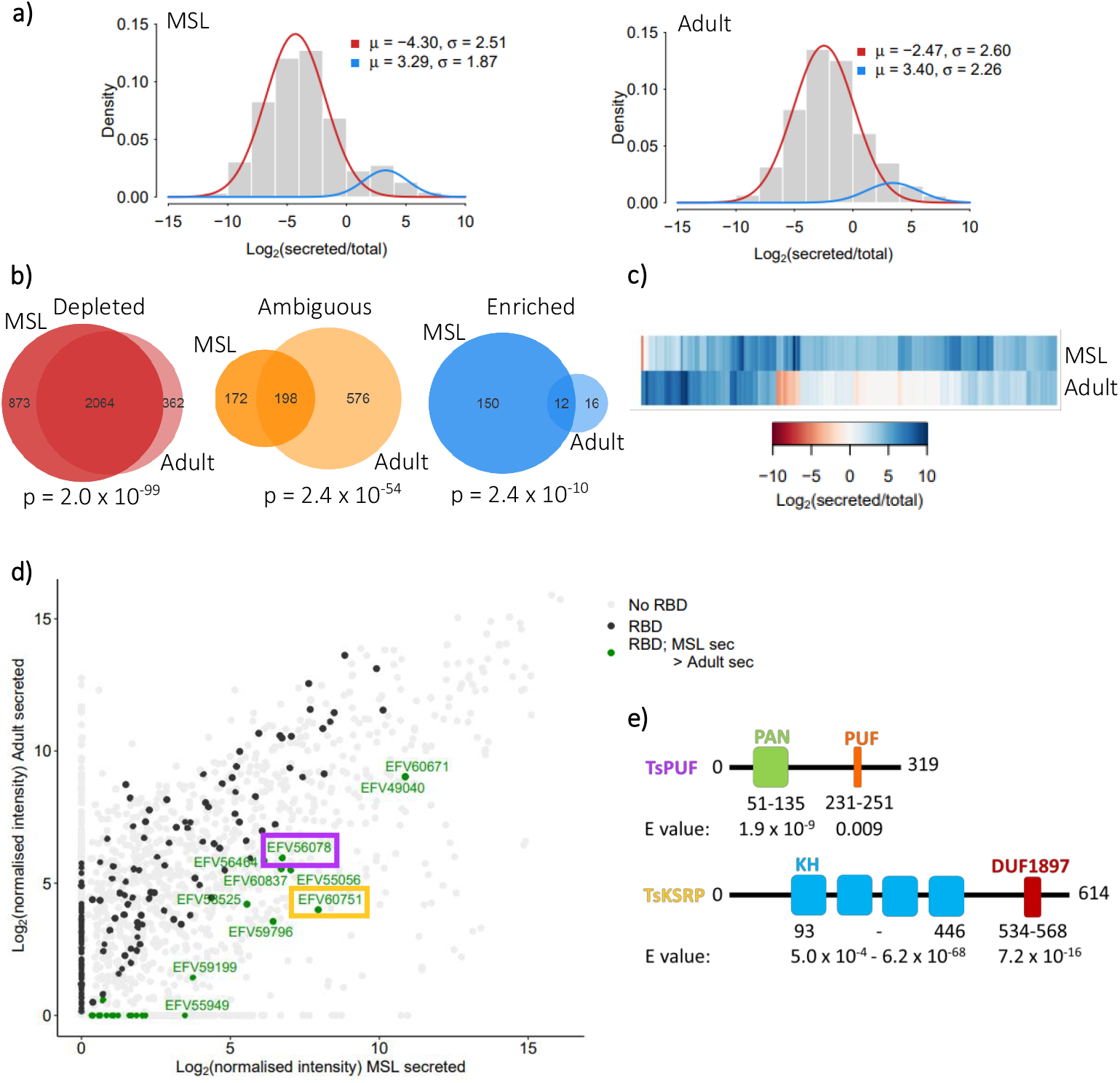
Comparison of *Trichinella spiralis* muscle-stage larvae (MSL) versus adult secretomes and identification of MSL abundantly secreted RNA-binding proteins. (a) Bimodal distribution of protein abundance in secreted material relative to in total worm in MSL and adults separately. Significant (p < 0.05) posterior probabilities from an expectation maximisation algorithm were used to define proteins as either enriched (blue) or depleted (red) in the secreted material. Proteins with no significant posterior probability were defined as ambiguous. (b) Venn diagrams comparing numbers of proteins enriched, depleted or ambiguous in secreted material in MSL versus adults. P values obtained from Fisher’s exact test of independence. (c) Heatmap comparing levels of enrichment of all 178 enriched secreted proteins in MSL versus adults. (d) Abundance, in MSL versus adults, of all proteins secreted by *T. spiralis.* Grey; no known RNA-binding domains (RBDs). Black; at least one RBD. Green; at least one RBD & more abundant in secreted material of MSL than that of adults (protein IDs labelled). Abundance of TsPUF (purple) and TsKSRP (orange) are highlighted. (e) Domain structure of the TsPUF and TsKSRP annotated by hmmscan. PUF; pumilio-*fem*-3 binding factor. KH; K homology domain. DUF1897; domain of unknown function 1897.

### Bioinformatic characterisation of TsPUF and TsKSRP

We characterised homologues of TsPUF across nematodes (Supplemental table 4), showing that TsPUF is widely conserved but that the region identified as the PUF domain was only evident within the *Trichinella* genus and a similar region within the *Trichuris* genus (Fig 2A,B). The N terminus of TsPUF contained a canonical signal peptide, followed by a Panhandle (PAN) domain (Fig 2C). The PAN domain is often found in extracellular proteins where it mediates protein-protein interactions[24]. These features suggested that TsPUF is most likely targeted to the extracellular environment through the canonical secretory pathway. Only one copy of the PUF domain was present (Fig 2C), in contrast to known Pumilio homology domain proteins where several tandem PUF domains are found with each domain responsible for contacting one nucleic acid on the RNA target[25,26]. The PUF region in TsPUF is short and not widely conserved so is likely to have convergently evolved similarity to the PUF repeat found in Pumilio family members. However, the presence of a sequence with similarity to the PUF repeat suggested that it might nevertheless have RNA binding properties. Consistently, an alpha-fold model for TsPUF predicted a folded structure with the PUF-like region on the surface of the protein (Fig 2D).

**Figure 2.**
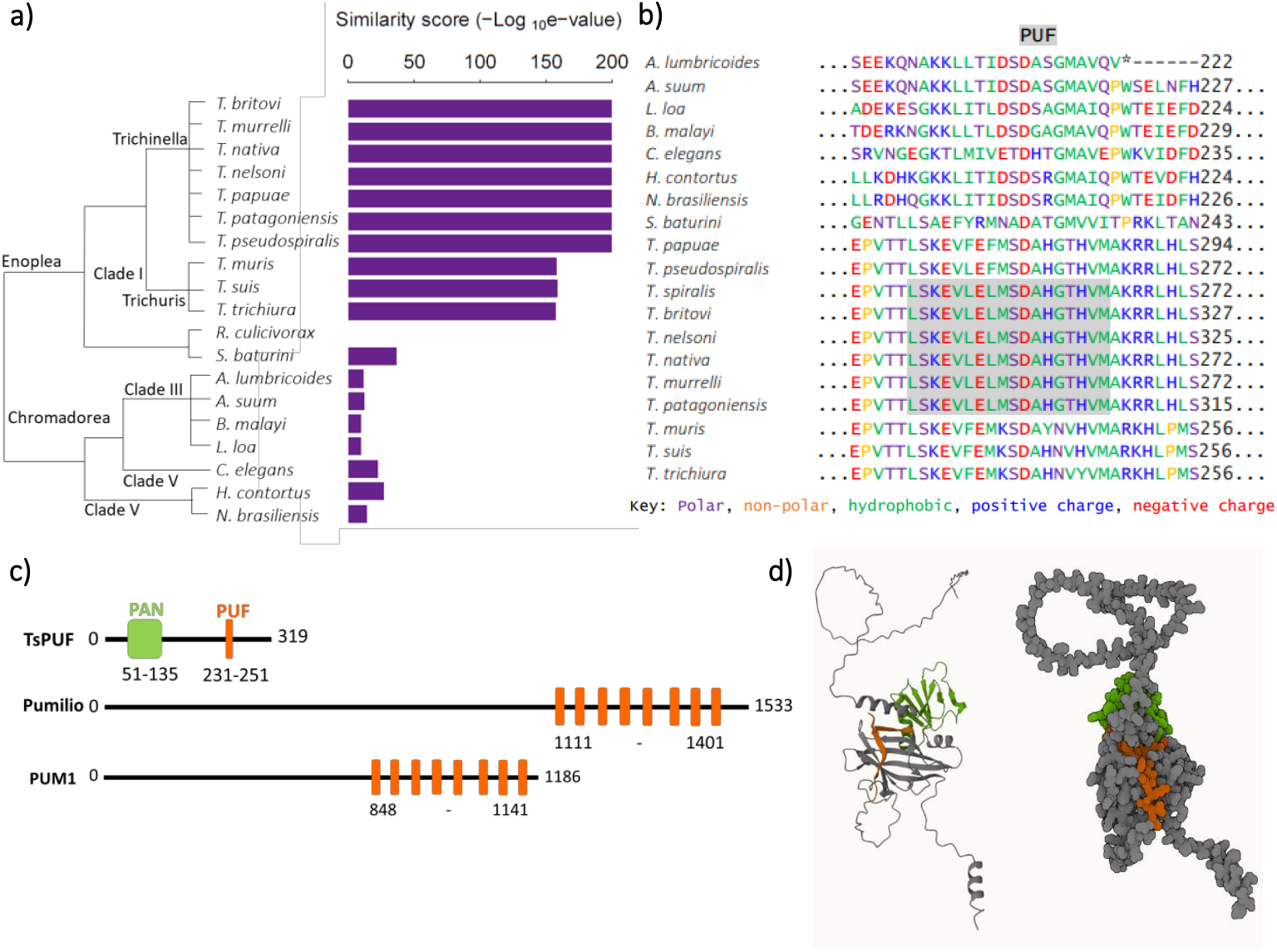
Characterisation of TsPUF protein and structure. (a) Similarity between TsPUF and its homologues in 19 nematode species; *Trichinella britovi, murrelli, nativa, nelsoni, papuae, patagoniensis* and *pseudospiralis, Trichuris muris, suis* and *trichiura, Romanomermis culicivorax, Soboliphyme baturini, Asacaris lumbricoides* and *suum, Brugia malayi, Loa loa, Caenorhabditis elegans, Haemonchus contortus* and *Nippostrongylus brasiliensis*. Homologues identified by performing a reciprocal best blast hit search. Similarity score = inverse log_10_ of the e value from the reciprocal best blast hit. (b) Alignment of the PUF region in TsPUF against the TsPUF nematode homologues. Multiple sequence alignment performed using Clustal Omega. Hmmscan used to identify Pfam domains. Amino acids are coloured according to their properties and the position of the PUF domain is highlighted. (c) Comparison of TsPUF protein domain structure versus that of two canonical PUF proteins; *Drosophila melanogaster* pumilio and *Homo sapiens* PUM1. (d) Alphafold prediction of TsPUF (A0A0V1BXK5_TRISP) structure. The PUF domain is coloured in orange and the PAN domain in green.

We next characterised TsKSRP using bioinformatics. TsKSRP was highly conserved across nematodes (Fig 3A). It was clearly homologous to characterised KSRP from other organisms, showing a similar domain structure to mammalian KSRP (Fig 3B). No signal peptide was present, nor extracellular domains. Alpha fold predicted a folded structure with the GXXG loop, previously implicated in nucleic acid binding[23] exposed to solvent (Fig 3C). This suggested that TsKSRP may have a similar function in intracellular RNA metabolism as in other organisms[27]. In the absence of a canonical signal peptide and with no domains typical of extracellular proteins, TsKSRP may be secreted from cells via alternative routes to the canonical secretory pathway (see discussion).

**Figure 3.**
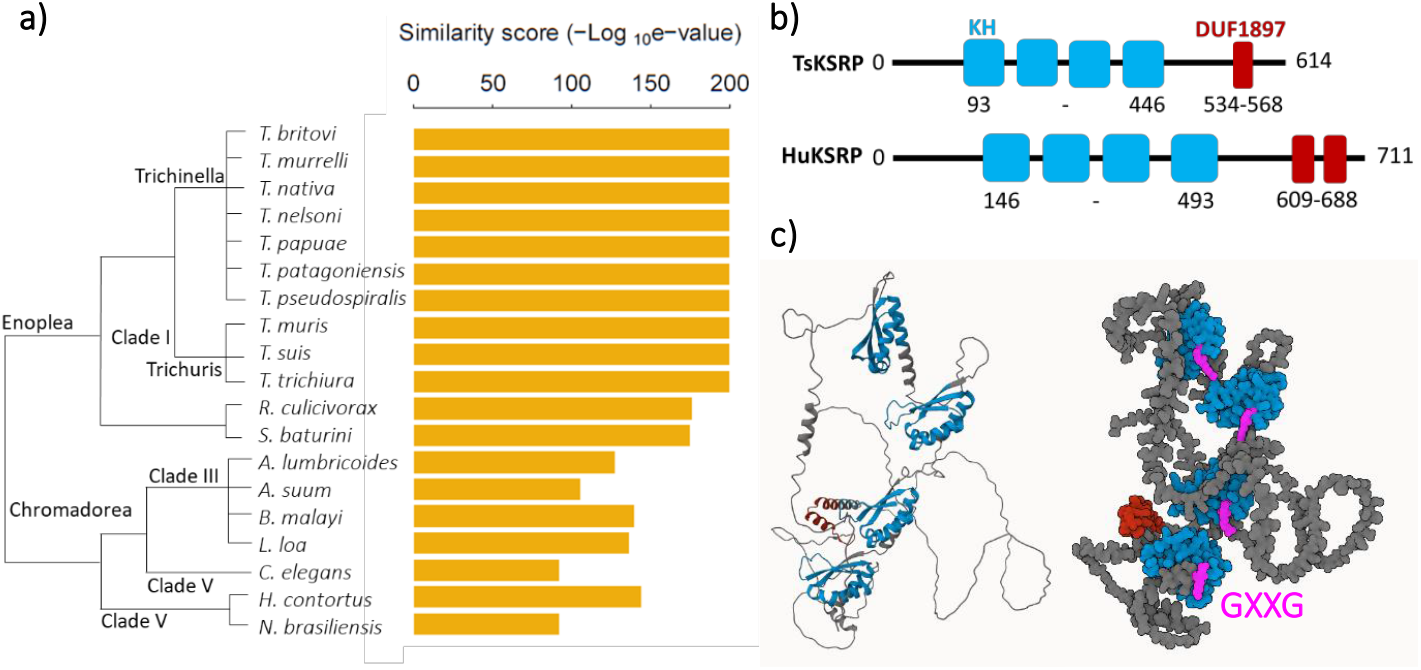
Characterisation of TsKSRP protein and structure. (a) Similarity between TsKSRP and its homologues in 19 nematode species; *Trichinella britovi, murrelli, nativa, nelsoni, papuae, patagoniensis* and *pseudospiralis, Trichuris muris, suis* and *trichiura, Romanomermis culicivorax, Soboliphyme baturini, Asacaris lumbricoides* and *suum, Brugia malayi, Loa loa, Caenorhabditis elegans, Haemonchus contortus* and *Nippostrongylus brasiliensis*. Homologues identified by performing a reciprocal best blast hit search. Similarity score = inverse log_10_ of the e value from the reciprocal best blast hit. (b) Comparison of TsKSRP protein domain structure versus that of human KSRP protein (HuKSRP). (c) Alphafold prediction of TsKSRP (A0A0V1B7I9_TRISP) structure. The KH domains are coloured in blue with the GXXG loop in pink. The DUF1897 domain is coloured in dark red.

### Recombinant TsKSRP and TsPUF bind miRNAs

To examine whether TsKSRP and TsPUF could contribute to the secretion of miRNAs by *T. spiralis* we tested whether TsKSRP and TsPUF could bind RNA in vitro. We expressed TsPUF with N-terminal his- and c-myc tags in yeast and purified from secreted material using Ni-NTA affinity chromatography. We expressed TsKSRP in bacteria with C-terminal his- and c-myc tags and purified using Ni-NTA affinity. As a positive control, we identified a *T. spiralis* Argonaute homologue (TsAGO) predicted to bind miRNAs[28], expressed it in bacteria with C-terminal his- and c-myc tags and purified it using Ni-NTA affinity. We incubated all three proteins with total RNA extracted from whole MSL, repurified the proteins using anti- c-myc pulldowns and extracted co-purifying RNA (Figure 4A). We then subjected co-purifying sRNAs to high-throughput sequencing, adding synthetic short non-coding RNAs with no overlap to the *T. spiralis* genome as normalization controls (see methods) (Supplemental table 5). The profile of reads in all reactions is visualised in Supplemental Figure 1. We focussed on miRNAs as our aim was to discover the mechanism of miRNA secretion and stability. However, we note that miRNAs make up a small percentage of the reads in all reactions and future work will be required to investigate whether other species of RNAs interact with these proteins.

**Figure 4.**
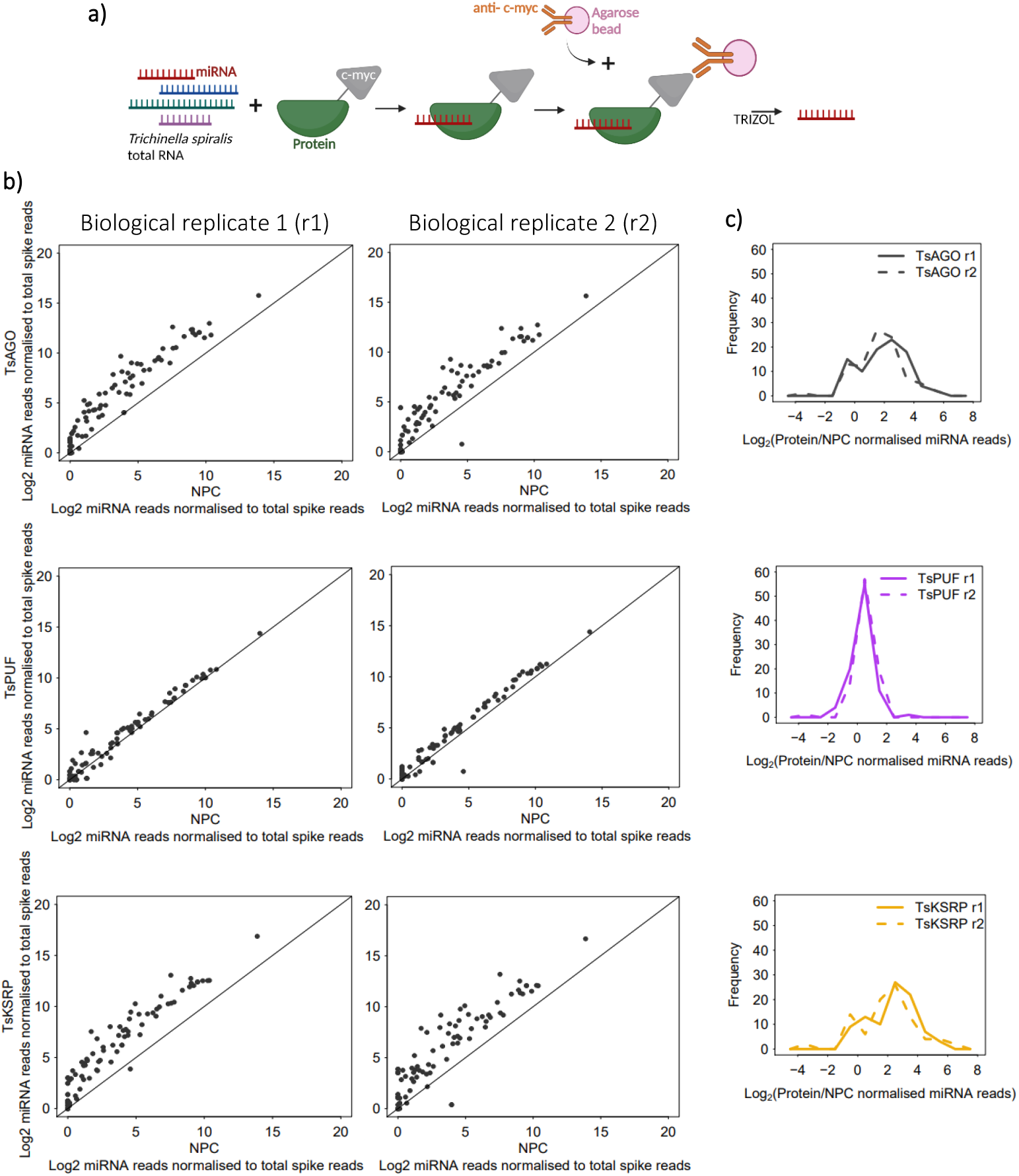
Analysis of miRNAs pulled down by *in vitro* RNA immunoprecipitation (RIP) using recombinant TsAGO/PUF/KSRP. (a) Schematic of RIP assay. (b) Enrichment of *T. spiralis* miRNA reads in two biological replicate (r) RIP reactions containing recombinant proteins (TsAGO/PUF/KSRP), relative to RIP control reactions with no protein (NPC). miRNA reads from sequencing of small RNA libraries were normalised against reads for oligos spiked in before sRNA library preparation. (c) Distribution of enrichment of *T. spiralis* miRNA reads in RIP reactions containing recombinant TsAGO/PUF/KSRP, relative to a NPC.

TsAGO and TsKSRP both bound selectively, with some miRNAs consistently enriched in the pulldown compared to others (Figure 4B and C). The distribution of enrichments was bimodal suggesting a small population of highly enriched miRNAs. There was a significant overlap of highly enriched miRNAs in the two biological replicates (Supplemental Figure 2). The enrichment of each miRNA was also highly correlated between TsKSRP and TsAGO pulldowns (Supplemental Figure 3). Human KSRP has some sequence-specific binding properties, in particular showing preference for G nucleotides in miRNAs[29] and discrimination against C nucleotides in all RNA targets[30]. We tested whether nucleotide content was different in miRNAs binding to TsKSRP or TsAGO in vitro. We found a significant depletion of C nucleotides in miRNAs enriched for TsKSRP binding but no enrichment for G content (Supplemental Figure 4). We did not find any significant enrichments for dinucleotides or trinucleotides (Supplemental Figures 5 and 6).

TsPUF exhibited a different pattern of enrichment from TsKSRP and TsAGO, whereby almost all miRNAs bound to a similar extent (Figure 4B and C). The unimodal distribution of enrichments thus suggested moderate, non-selective binding to most miRNAs (Figure 4B). Furthermore, the correlations of enrichments between TsPUF and either TsKSRP or TsAGO were weak (Supplemental Figure 3), supporting a different binding mode. We wondered whether the PUF-like region in TsPUF contributed to RNA binding, so we expressed and purified recombinant TsPUF lacking specifically this region (Figure 5A and B). TsPUF lacking the PUF-like region failed to bind miRNA, suggesting that despite lack of homology to canonical PUF proteins, the PUF-like region may contribute to RNA binding either directly or through stabilising the correct fold of the protein (Figure 5C and D).

**Figure 5.**
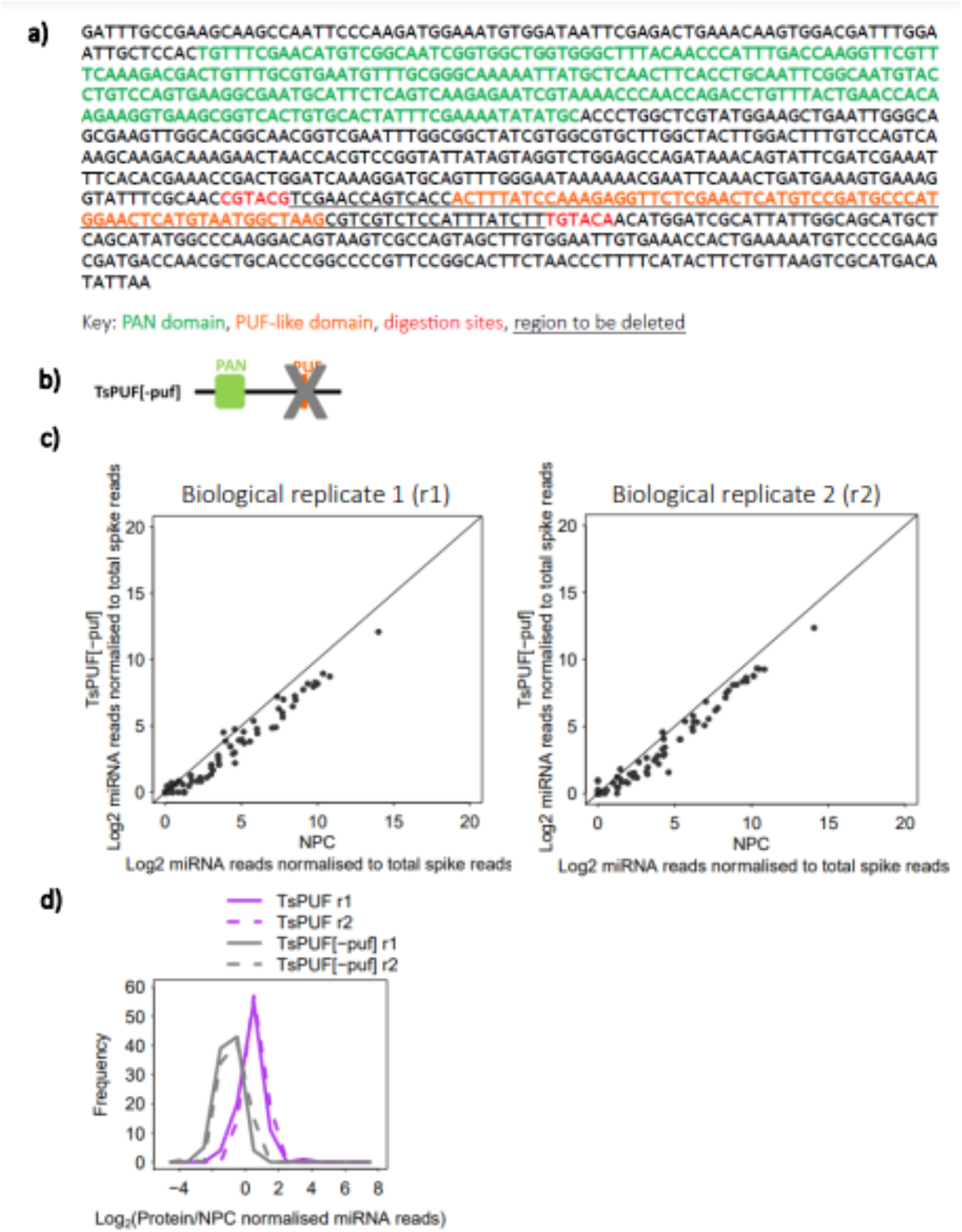
Analysis of miRNAs pulled down by *in vitro* RNA immunoprecipitation (RIP) using recombinant TsPUF mutant without PUF domain (TsPUF[-puf]). (a) Compatible digestion sites BsiWI (CGTACG) and BsrGI (TGTACA) were used to cut-out the region containing the PUF-like domain from the TsPUF gene. Only the protein-coding sequence (post-signal peptide and tag sequences) are shown. (b) Domain structure of TsPUF[-puf] protein. (c) Enrichment of *T. spiralis* miRNA reads in two biological replicate (r) RIP reactions containing TsPUF[-puf], relative to RIP control reactions with no protein (NPC). miRNA reads from sequencing of small RNA libraries were normalised against reads for oligos spiked in before sRNA library preparation. (d) Distribution of enrichment of *T. spiralis* miRNA reads in RIP reactions containing recombinant TsPUF/TsPUF[-puf], relative to a NPC.

Taken together we concluded that the secreted proteins TsKSRP and TsPUF bind miRNAs, but whilst TsKSRP showed selective binding, similar to the canonical sRNA binding protein TsAGO, TsPUF binds non-selectively to miRNAs.

### TsPUF binds miRNAs in the secreted material from T. spiralis larvae

Having established that recombinant TsKSRP and TsPUF bound to miRNAs in vitro, we next wanted to test whether the proteins were present bound to RNA in the secretome of *T. spiralis*. We were not able to raise a specific antibody against TsKSRP. However, we successfully raised an antibody against TsPUF which produced a single, clear band at the correct size when tested by western blot on secreted material, indicating specific binding (Figure 6A). We therefore focussed on TsPUF for this analysis. We visualised the localisation of TsPUF on fixed sections from muscle of infected mice using immunofluorescence. We observed highly specific staining for TsPUF in larvae (Figure 6B). The protein had an extracellular localisation, most strongly concentrated in the pseudocoelomic fluid (Figure 6B). This was consistent with the prediction that TsPUF is targeted to the conventional secretory pathway. We did not observe any TsPUF in mouse muscle cytoplasm; however, this does not exclude secretion from the parasite into the host muscle cells, as secreted proteins may be too diffuse in host tissue to be detected.

**Figure 6.**
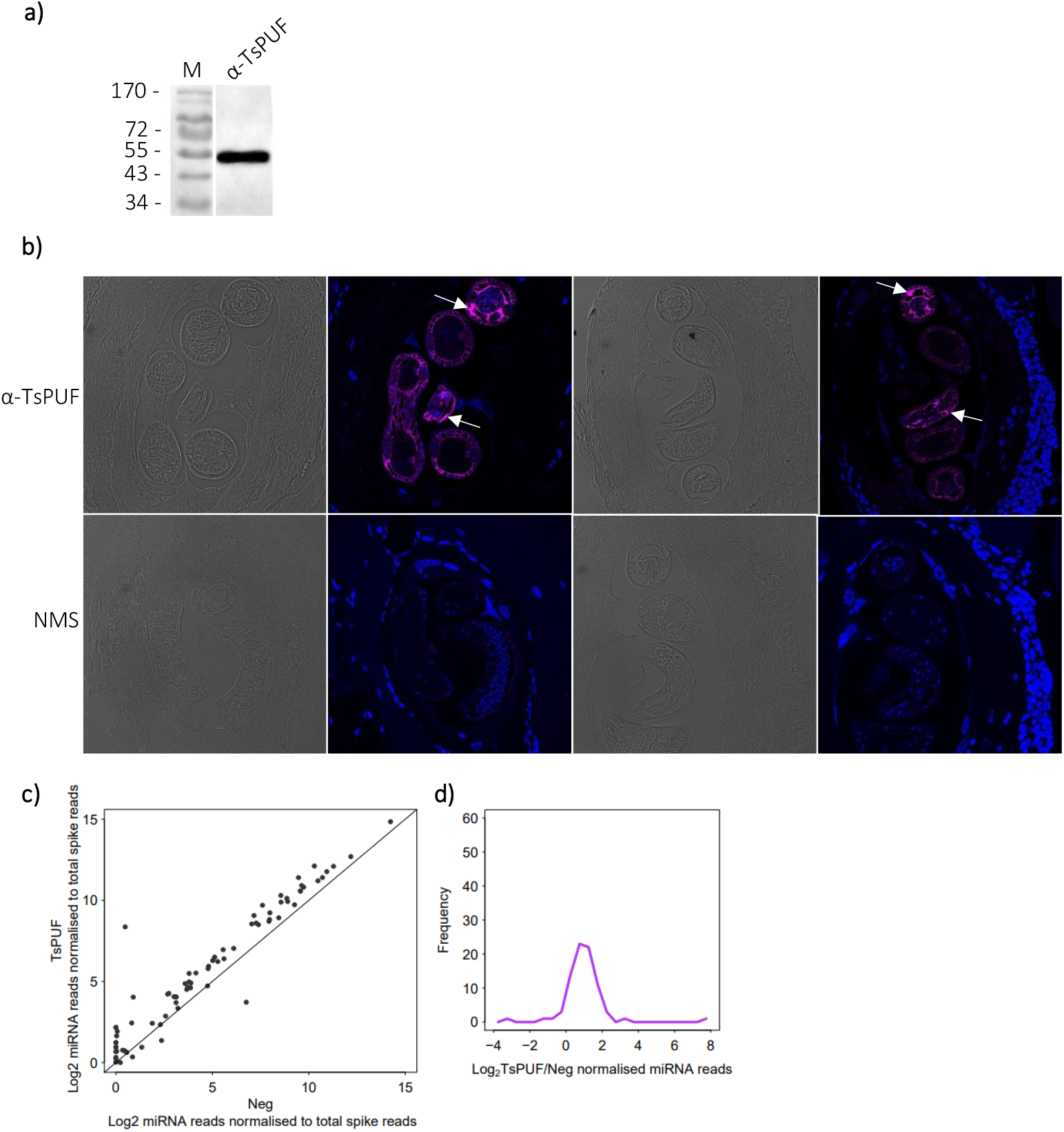
*In vivo* characterisation of native TsPUF in *Trichinella spiralis* muscle-stage larvae (MSL). (a) Western blot performed on *T. spiralis* MSL secreted material using α-TsPUF antiserum. M = marker, with molecular weight (kDa) indicated. (b) Immunofluorescence staining of *T. spiralis* infected mouse thigh muscle sections. Top panels stained with α-TsPUF antiserum and DAPI counterstain (brightfield and immunofluorescence), with localisation in larval pseudocoelom arrowed. Bottom panels stained with naïve mouse serum (NMS) and DAPI counterstain (brightfield and immunofluorescence). (c) Enrichment of *T. spiralis* miRNA reads by RNA immunoprecipitation (RIP) of native TsPUF, using α-TsPUF antiserum, from MSL secreted material. Enrichment is relative to a RIP control reaction with naïve mouse serum (Neg). miRNA reads from sequencing of small RNA libraries were normalised against reads for oligos spiked in before sRNA library preparation. (d) Distribution of enrichment of *T. spiralis* miRNA reads by RIP of native TsPUF, relative to Neg.

We next tested whether the protein was bound to miRNAs in secreted material collected from *T. spiralis* MSL in culture. We immunoprecipitated (IP) TsPUF from secreted material and extracted bound RNAs, comparing to IP with naïve mouse serum as a control (Supplemental table 6). TsPUF IPs showed clear enrichment of most miRNAs compared to the negative control (Figure 6C), consistent with relatively non-selective binding. There was little correlation between the enrichment of miRNAs bound to recombinant TsPUF and immunoprecipitated TsPUF (Supplemental Figure 7). Thus we concluded that TsPUF binds to miRNAs in the *T. spiralis* MSL secretome non-selectively. Future work would benefit from investigating whether TsPUF is bound to other species of RNA in the MSL secretome. The profile of reads in the pulldown is visualised in Supplemental Figure 8.

## Discussion

Here, using a combination of computational biology, biochemistry and cell biology, we discovered two RNA binding proteins secreted by the parasitic nematode *T. spiralis*. We confirmed that recombinant versions of both proteins bind miRNAs in vitro. In the case of one of these proteins, TsPUF, we were able to demonstrate that it bound miRNAs in secreted material from the parasite. Our findings provide new insights into the mechanism whereby miRNAs might be secreted and stabilised in parasitic nematodes and may be relevant for understanding the biology of extracellular RNA more widely.

### Secreted RNA binding proteins in T. spiralis

We characterised two RNA binding proteins that were present in secreted material from *T. spiralis*. One of these proteins, TsKSRP, was a member of a family of known RNA binding proteins. KSRP in other organisms is associated with miRNA sorting and stabilisation[27], and we showed that TsKSRP has selective binding to some miRNAs. However, it has never been characterised as an extracellular protein. This raises the question of how KSRP is secreted by *T. spiralis*. Importantly, we did not detect a canonical signal peptide in the protein sequence, suggesting that it may not be secreted via the ER secretory pathway. One possibility is that the protein is secreted via exosomes which are prone to lysis, releasing KSRP into extravesicular material.

In contrast, TsPUF contained the PAN domain, often found in extracellular proteins, and had a signal peptide, thus is likely to be secreted via the ER. Identification of this protein as an RNA binding protein was due to a small region that showed weak similarity to a PUF repeat; however, we showed that this is unlikely to reflect homology to the Pumilio family and may be either convergent evolution or a coincidence. Although a deletion encompassing the PUF repeat failed to bind RNA we cannot exclude that this interfered with the overall structure. Interestingly, it showed a very different profile of RNA binding to either TsKSRP or TsAGO, suggesting an entirely different mechanism of RNA binding and stabilisation. Overall, TsPUF is a novel RNA binding protein but the exact mechanism responsible for RNA binding still awaits characterisation. We note that extracellular miRNAs are very common across organisms but there is so far limited evidence for canonical RNA binding proteins involved in their stabilisation[9]; indeed novel RNA binding proteins in extracellular material have been uncovered[31]. It is possible that our discovery of TsPUF as an RNA binding protein through domain searches was serendipitous and we predict that there may be many other non-canonical RNA binding proteins with roles in RNA secretion or stabilisation outside cells.

### Insights into the mechanism of RNA secretion by T. spiralis

The different properties of KSRP and TsPUF enable us to speculate on the pathway of RNA secretion from *T. spiralis* MSL. KSRP in humans is able to bind to a variety of RNA targets[32], including miRNA precursors via an interaction with the unpaired region of the stem-loop[29]. This interaction is proposed to underpin the requirement for KSRP in processing specific miRNAs[29]. KSRP has not been reported as binding to mature miRNAs, but given its propensity for single strand RNA binding it is possible that it could interact with them directly, potentially following Dicer cleavage. Human KSRP shows a preference for binding G-rich miRNAs and selects against cytosine nucleotides[29,30]. We found that TsKSRP binds selectively to miRNAs and that these miRNAs were depleted of cytosine, although we did not detect enrichment of G within enriched miRNAs. It is therefore possible that TsKSRP aids selective export of miRNAs, utilising sequence-specific binding. For this to be feasible, folded TsKSRP in complex with miRNA would have to be able to move into the extracellular space. Exactly how this could occur is unclear but suggestions could include a vesicle which lyses after secretion or a yet undiscovered direct route for folded proteins through the plasma membrane[10]. In contrast, TsPUF is most likely to be secreted in an unfolded form through the canonical protein secretion pathway. It would thus only be able to bind to miRNAs once it reaches the extracellular space. We therefore propose that KSRP transfers miRNAs to TsPUF via a hand-off mechanism. As a non-selective binder, TsPUF would therefore acts as a “sponge” to stabilise all extracellular RNAs. This may enable delivery of extracellular RNAs to host complexes inside infective cells, or TsPUF-miRNA complexes may be able to target host genes directly. It will be intriguing to test the extent to which this mechanism operates in muscle cells infected with *T. spiralis* and what role this might play in its pathogenesis.

## Methods

### Isolation of T. spiralis adults and MSL and preparation of total worm extract, secreted material and total worm RNA

Adult parasites were collected from infected rat intestines 6 days post-infection by sedimentation in a Baermann funnel and MSL were recovered from digested mouse muscle 2 months post-infection, as previously described[33]. For preparation of total worm extracts, adults/MSL were lysed in PBS with 0.05% Tween 20 and protease inhibitors using a Qiagen TissueLyser II. The lysate was collected as the total worm extract. For preparation of secreted material, parasites were cultured in serum-free medium for up to 72 hr as previously described[33]. Secreted products were collected daily and the supernatants cleared through 0.2 μm filters, pooled and concentrated using 10000 molecular weight cutoff vivaspin columns. For preparation of total worm RNA, parasites were lysed in TRIZOL using the TissueLyser followed by standard TRIZOL manufacturer’s guidelines and RNA precipitation.

### Analysis of T. spiralis proteins by mass spectrometry and identification of RNA-binding candidates

Matrix assisted laser desorption ionisation time of flight (MALDI-TOF) mass spectrometry was performed on total worm extracts and secreted material from *T. spiralis* MSL and adults (two technical replicates each). All figures and analysis of the proteomic datasets were performed in RStudio. Protein abundance was defined as the mean normalised intensity value from mass spectrometry. To identify potential RNA-binding proteins in the MSL secretome, all secreted proteins were searched for RNA-binding domains. First, a literature search was performed to create a list of 59 canonical/non-canonical RNA-binding domains[34–36] (Supplemental table 2). Next, hmm-scan in the HMMER software (hmmer.org) was used to search the amino acid sequences of all secreted proteins for all domains in the Pfam database[35]. The identified domains were then cross-matched with the curated list of RNA-binding domains.

### Conservation and bioinformatic characterisation of TsPUF and TsKSRP

The proteomes of *T. spiralis* and other nematode species were downloaded from Wormbase Parasite [37] (Supplemental table 4). A Blast search[38] was then performed for TsPUF (EFV56078) and TsKSRP (EFV60751) to find hits in each nematode proteome. The reciprocal search was then performed; a Blast search of the hits in each nematode against the *T. spiralis* proteome. The reciprocal best blast hit was defined as a homologue. Clustal Omega[39] was used to perform multiple sequence alignments of the homologues and hmm-scan used to identify Pfam domains. Alphafold[40,41] was used to predict the model structure of TsPUF (A0A0V1BXK5_TRISP) and TsKSRP (A0A0V1B7I9_TRISP). Mol*Viewer[42] was used to produce model images from the alphafold PDB files.

### Recombinant protein expression in bacteria

Total RNA extracted from *T. spiralis* muscle-stage larvae was reverse transcribed (RT) using Superscript IV reverse transcriptase (ThermoFisher) following the manufacturer’s protocol. A Q5 high fidelity PCR reaction (New England Biolabs) was performed on the cDNA to amplify TsKSRP and TsAGO genes using primers encoding a c-myc tag at the 3’ end. The genes were ligated with a pET21a(+) vector, encoding a his-tag, via compatible restriction digestion sites. Plasmids were replicated and isolated from *Escherichia coli* DH5α for transformation into the expression host *E. coli* BL21. TsKSRP and TsAGO were batch produced via large scale culture of transformed BL21 and induction of protein expression using 1 mM IPTG at 18°C overnight. Cells were lysed with 2.5 mg/ml lysozyme via 3x freeze-thaw cycles, incubation on ice for 2 hr and sonication. Ni-NTA affinity chromatography was performed to purify his-tagged TsAGO and TsKSRP from lysates, following the standard QIAexpressionist protocol.

### Recombinant secretory protein expression in yeast

A TsPUF gene, with sequences encoding a his- and c-myc tag at the N-terminus, was synthesised by GeneArt (ThermoFisher). The TsPUF mutant without the PUF domain (TsPUF[- puf]) was created by digesting the TsPUF gene with restriction enzymes that cut in positions either side of the sequence encoding the PUF domain and ligating it back together. Both TsPUF[-puf] and the wildtype TsPUF were ligated with a pPICZα vector via compatible restriction digestion sites. Plasmids were replicated and isolated from *E. coli* DH5α for transformation into the expression host *Pichia pastoris*. TsPUF and TsPUF[-puf] were batch produced via large scale culture of transformed *P. pastoris* and induction of protein expression using methanol. Secreted recombinant proteins were collected from the supernatant of the yeast culture. Ni-NTA affinity chromatography was performed to purify his-tagged TsPUF and TsPUF[-puf] from the supernatant, following the standard QIAexpressionist protocol.

### RNA immunoprecipitation using recombinant proteins

Recombinant protein (347 nM) was incubated with total RNA (156 nM) extracted from *T. spiralis* MSL for 30 min in PBS with 0.05% Tween 20 (PBST). The protein:RNA mixture was then incubated with anti- c-myc agarose beads from a Pierce c-Myc Tag IP/Co-IP kit (ThermoFisher) for 2.5 hr. The protein:RNA:anti-c-myc mixture was washed through Pierce spin columns with PBST to remove anything unbound to the anti- c-myc agarose beads. TRIZOL was then used to elute the protein:RNA from the beads. Protein:RNA in TRIZOL was disrupted by repeated freeze-thawing in liquid nitrogen and vigorous vortexing. Chloroform was used to allow phase separation and subsequent precipitation of RNA in the aqueous phase with glycogen and isopropanol. The RNA pellet was washed with 75% EtOH and resuspended in ultrapure water. No protein control (NPC) reactions were performed in the exact same way, except with no recombinant protein incubated with the RNA.

### Preparation of cDNA libraries for small RNA sequencing

RNA was used to prepare cDNA libraries with a Truseq sRNA library prep kit (Illumina), following the manufacturer’s protocol. Single end sequencing of libraries was kindly performed by the MRC LMS Genomics Laboratory on a NextSeq2000 machine. Oligos were spiked in at the library preparation stage at a constant concentration (0.2% of the input RNA) to allow normalisation of reads from different libraries. To identify appropriate oligos, all human miRNA sequences were downloaded from miRbase[43] and then a Blast search was performed to identify those with no match against the *T. spiralis* genome. Three sequences were selected in order to have a range of sequence lengths from 19-23 nucleotides long (GGCUUGCAUGGGGGACUGG, UGACAGCGCCCUGCCUGGCUC and GUUUGCACGGGUGGGCCUUGUCU). Oligos were synthesised by Merck.

### Bioinformatic analysis of small RNA sequencing data

Sequencing reads were demultiplexed by the MRC LMS Genomics Laboratory. A shell script was then used to trim adapters and convert to collapsed fasta files with the ‘fastx’ package. To identify *T. spiralis* miRNA, bowtie was used to find the position in the *T. spiralis* genome where the reads align. Bedtools[44] was then used to find reads that align with our previous annotation of miRNAs in the *T. spiralis* genome[28]. All analyses were then performed in RStudio. Information on the first nucleotide and length for each read was extracted using a custom Perl script. For analysis of single nucleotide occurrence in the miRNA sequences, the proportion of each sequence made up of each nucleotide was calculated. The mean nucleotide occurrences were then calculated for miRNA sequences in different enrichment groups. For analysis of di-/tri-nucleotide occurrence in the miRNA sequences, the occurrence of all possible di-/tri-nucleotides were counted in each sequence. These counts were then normalised against the total number of di-/tri-nucleotides present in the given sequence. The mean normalised di-/tri-nucleotide occurrences were then calculated for miRNA sequences in different enrichment groups. To look for significant single/di-/tri-nucleotides, Chi Squared tests, followed by Bonferroni correction, were performed on the mean occurrences of relevant single/di-/tri-nucleotides in different enrichment groups.

### Generation of TsPUF antiserum in a mouse

Endotoxins were removed from purified recombinant TsPUF using Pierce Endotoxin Removal Resin Columns, following the manufacturer’s protocol. A mouse was immunised with the endotoxin-free protein mixed with Imject Alum (Thermo Scientific) as an adjuvant. The mouse was then boosted with more protein/adjuvant mix 4 weeks later, and twice more 2 weeks apart before bleeding the mouse 1 week after the final boost. The supernatant was then collected from the blood and this was used as the TsPUF antiserum.

### Immunofluorescence on infected muscle tissue to visualise TsPUF

A section of thigh muscle from an infected mouse was fixed in 10% neutral buffered formalin, embedded in paraffin and sectioned using standard techniques. The sections were deparaffinized and rehydrated via a series of washes with xylene and decreasing concentrations of ethanol. The sections were blocked with 1% bovine serum albumin (BSA) in PBS with 0.1% Tween 20 (PBST) and 10% naïve goat serum. A second block was performed (for mouse on mouse staining) using goat F(ab) anti-mouse IgG (Abcam) at 1:100 in PBST. Sections were then incubated with TsPUF antiserum (or naïve mouse serum) at 1:200 in PBST with 1% BSA. Goat anti-mouse alexa fluor 488 (Abcam) at 1:500 in PBST with 1% BSA was then used to stain the sections. The sections were then counterstained using mounting medium containing DAPI (Abcam). Immunofluorescence staining was visualised using a Leica SP8 - STELLARIS 5 Inverted Light Sheet Confocal Microscope. Localisation was determined by reference to the structure of infective larvae[45].

### RNA immunoprecipitation of native TsPUF from MSL secreted material

Immunoprecipitations were performed with 10 ul TsPUF antiserum (or 10 ul naïve mouse serum), incubated with 100 ug (protein concentration) *T. spiralis* MSL secreted material in PBS with 0.05% Tween 20 (PBST) for 1 hr. The antiserum:secreted material mixture was then incubated with 50 ul Dynabeads Protein G magnetic beads (ThermoFisher) for 30 min. A magnetic plate was used for washing with PBST. TRIZOL was then used to elute the protein:RNA from the beads. RNA was then isolated following the same method used in the recombinant protein RNA immunoprecipitations.

## Acknowledgements

We thank Lorraine Lawrence (NHLI, Imperial College London) for processing tissue samples for histology.

## Ethics Statement

The animal study was reviewed and approved by the Animal Welfare Ethical Review Board at Imperial College London and was licensed by and performed under the UK Home Office Animals (Scientific Procedures) Act Personal Project Licence number 70/8193: ‘Immunomodulation by Helminth Parasites’.

## Data availability

Raw sequencing data has been deposited to the SRA and can be accessed via https://dataview.ncbi.nlm.nih.gov/object/PRJNA967312?reviewer=b29l2lhaompviqrpiolvq3 s453

Processed data

All scripts required for producing the figures have been posted to GitHub https://github.com/SarkiesLab/TspirExRNAProts

## Supplementary Figure Legends

Supplementary Figure 1. Profile of sequencing reads, in terms of length (in nucleotides; nt) and nt in the first position, of reads from *in vitro* RNA immunoprecipitation (RIP) reactions. RIP read profiles represent the mean of two biological replicates. Profile of reads from sequencing of total RNA extracted from *Trichinella spiralis* muscle-stage larvae is also shown (Total).

Supplementary Figure 2. Comparison of miRNAs pulled down by *in vitro* RNA immunoprecipitation (RIP) in two biological replicates (r). (a) Enrichment of *T. spiralis* miRNA reads in protein RIPs, relative to no protein control (NPC) RIPs, in r1 versus r2. (b) Distribution of enrichment of *T. spiralis* miRNA reads in protein RIP reactions. Thresholds to define the level of enrichment (in terms of log_2_(Protein/NPC)) are labelled in green and dark green. (c) Venn diagrams to show the overlap between miRNAs enriched in the two replicates. P values from Fisher’s exact test of independence.

Supplementary Figure 3. Comparison of the level of enrichment, relative to no protein control (NPC), of miRNAs by *in vitro* RNA immunoprecipitation (RIP) with different recombinant proteins. Enrichment values represent the mean of two biological replicate RIP reactions.

Supplementary Figure 4. Global nucleotide (a) and AU dinucleotide (b) abundance in miRNAs (very) enriched, versus not (very) enriched, by recombinant TsKSRP in RNA immunoprecipitation (RIP) reactions. Definitions of miRNA enrichment, relative to a RIP control reaction with no protein (NPC), are shown. (a) For every enrichment group, the proportion (%) of each sequence made up of each nucleotide was calculated. Values shown here are the average (mean) of these proportions within the group. Chi Squared tests, followed by Bonferroni correction, were performed on the mean values for every nucleotide to compare the following groups: Enr versus Not Enr, V.Enr versus Not Enr, V.Enr versus Not V.Enr. (b) For every enrichment group, the occurrence of AU in each sequence was counted and normalised against the total number of all dinucleotides in that sequence. Values shown are the mean normalised occurrences. Chi Squared tests were performed on the mean values to compare the following groups: enriched versus not enriched, enriched versus very enriched, enriched versus enriched shuffled, very enriched versus not enriched, very enriched versus not very enriched and very enriched versus very enriched shuffled. Only the differences with significant (<0.1) p values are labelled (*).

Supplementary Figure 5. Global dinucleotide abundance in miRNAs (very) enriched, versus not enriched/depleted, by recombinant TsAGO(a)/TsPUF(b)/TsKSRP(c) in RNA immunoprecipitation (RIP) reactions. For every enrichment group, the occurrence of each dinucleotide in each sequence was counted and normalised against the total number of all dinucleotides in that sequence. Values shown are the mean normalised occurrences. Chi Squared tests, and Bonferroni correction, were performed on the mean values for every dinucleotide to compare the following groups: very enriched versus not enriched and very enriched versus very enriched sequences shuffled x1000 (for TsAGO/TsKSRP). For TsPUF, enriched versus depleted and enriched versus enriched sequences shuffled x1000 were compared. No significant differences were found in any group.

Supplementary Figure 6. Global trinucleotide abundance in miRNAs (very) enriched, versus not enriched/depleted, by recombinant TsAGO(a)/TsPUF(b)/TsKSRP(c) in RNA immunoprecipitation (RIP) reactions. For every enrichment group, the occurrence of each trinucleotide in each sequence was counted and normalised against the total number of all trinucleotides in that sequence. Values shown are the mean normalised occurrences. Chi Squared tests, and Bonferroni correction, were performed on the mean values for every trinucleotide to compare the following groups: very enriched versus not enriched and very enriched versus very enriched sequences shuffled x1000 (for TsAGO/TsKSRP). For TsPUF, enriched versus depleted and enriched versus enriched sequences shuffled x1000 were compared. No significant differences were found in any group.

Supplementary Figure 7. Comparison of the level of enrichment of miRNAs by *in vitro* RNA immunoprecipitation (RIP) using recombinant TsPUF versus the level of enrichment by RIP of native TsPUF. Enrichment values are relative to a negative control (log_2_(protein/negative normalised miRNA reads)). The negative control is either a RIP reaction with no protein (for recombinant protein RIPs) or a RIP using naïve mouse serum (for native RIPs with anti-serum against the native proteins). Enrichment values for the recombinant RIP represents the mean of two biological replicate RIP reactions.

Supplementary Figure 8. Profile of sequencing reads, in terms of length (in nucleotides; nt) and nt in the first position, of reads from RNA immunoprecipitation (RIP) of native TsPUF from *Trichinella spiralis* muscle-stage larvae (MSL) secreted material. Profile of reads from sequencing of RNA extracted from *T. spiralis* MSL secreted material is also shown.

## Legends for Supplemental Tables

**Supplemental table 1.** Raw and processed data for the intensity level of all proteins identified by mass spectrometry in *Trichinella spiralis* muscle-stage larvae (MSL) and adult secreted material and total worm extracts.

**Supplemental table 2.** List of RNA-binding domains (RBDs) used in this study. List created by performing a literature search for canonical/non-canonical RBDs.

**Supplemental table 3.** Candidate RNA-binding domain-containing proteins which were more enriched in the secreted material of *Trichinella spiralis* muscle-stage larvae (MSL) than by the secreted material of adults. Abundance of all proteins is also shown (mean normalised intensity, from mass spectrometry, of two replicates).

**Supplemental table 4.** Nematodes used in this study to analyse the conservation of *Trichinella spiralis* proteins.

**Supplemental table 5.** *Trichinella spiralis* miRNA read counts, normalised to total oligo spike counts, from small RNA sequencing of RNA immunoprecipitated using recombinant TsKSRP, TsAGO, TsPUF and TsPUF[-puf]. Data for no protein controls (NPC) are also shown. Two biological replicates performed.

**Supplemental table 6.** *Trichinella spiralis* miRNA reads, normalised to total oligo spike counts, from small RNA sequencing of RNA immunoprecipitated from *T. spiralis* secreted material in a native TsPUF pull-down. Data for a negative control (Neg) is also shown.

