## Supplementary Figures for "Identification of proteins that bind extracellular microRNAs secreted by the parasitic nematode *Trichinella spiralis*"

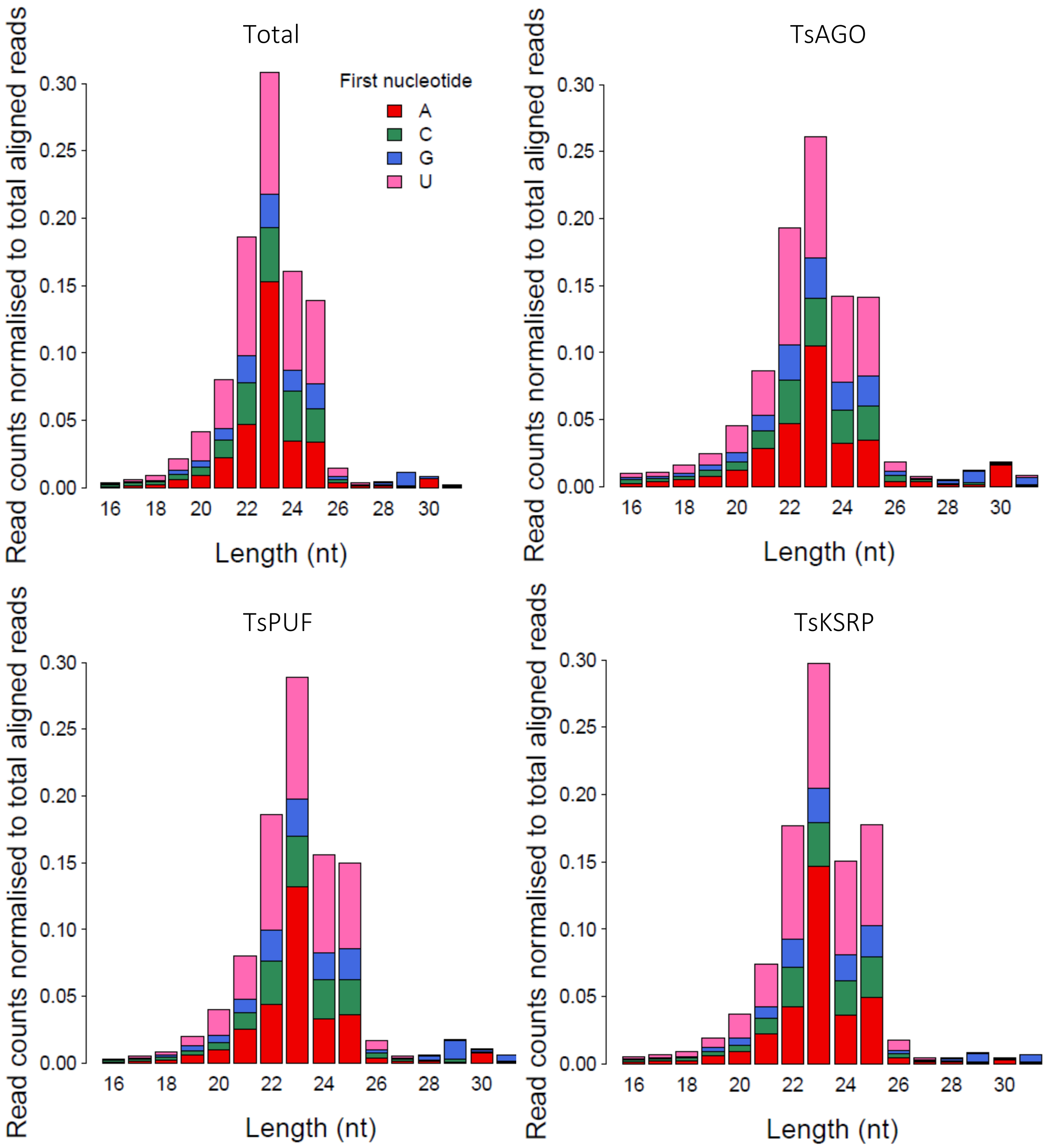


Supplementary Figure 1.

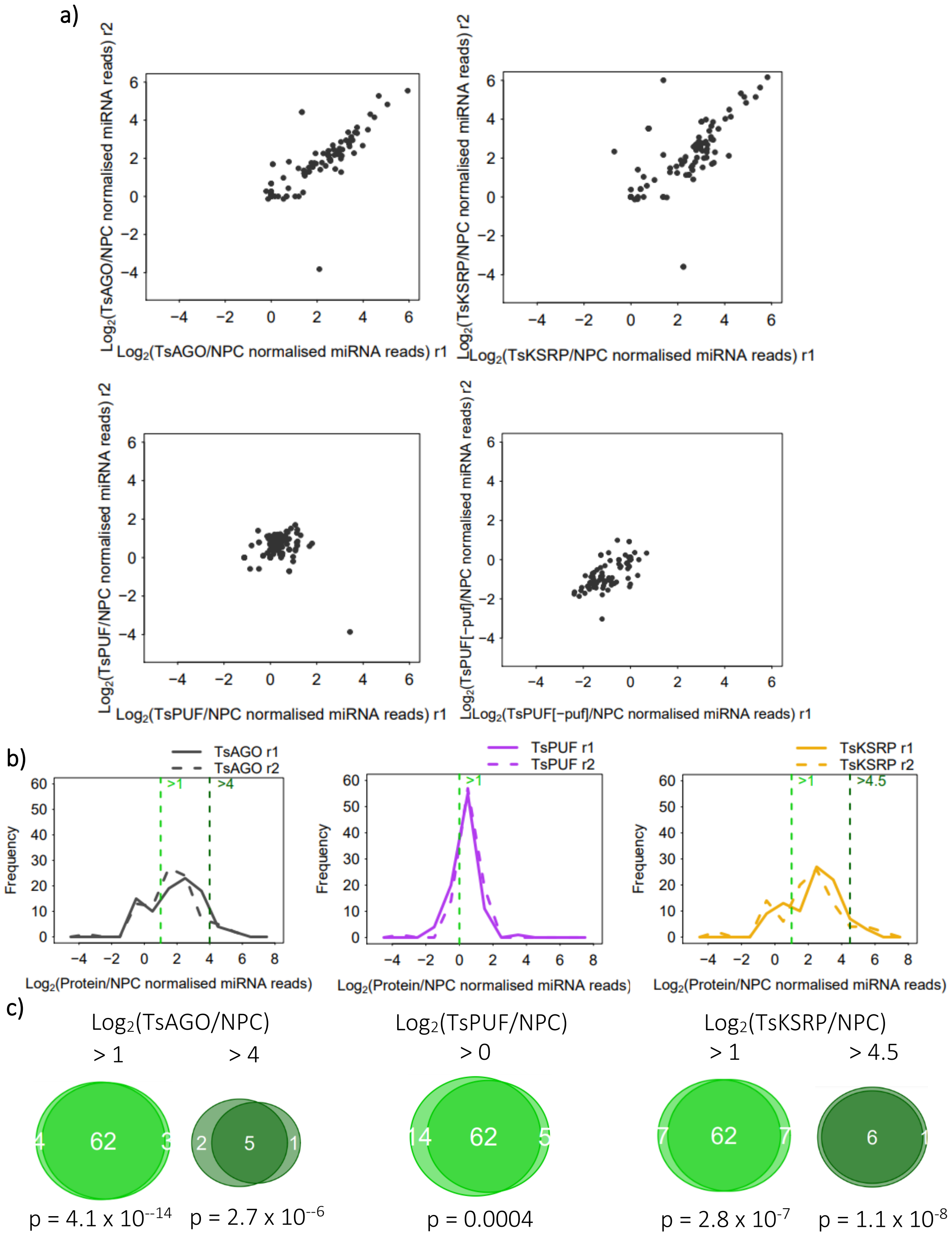


Supplementary Figure 2.

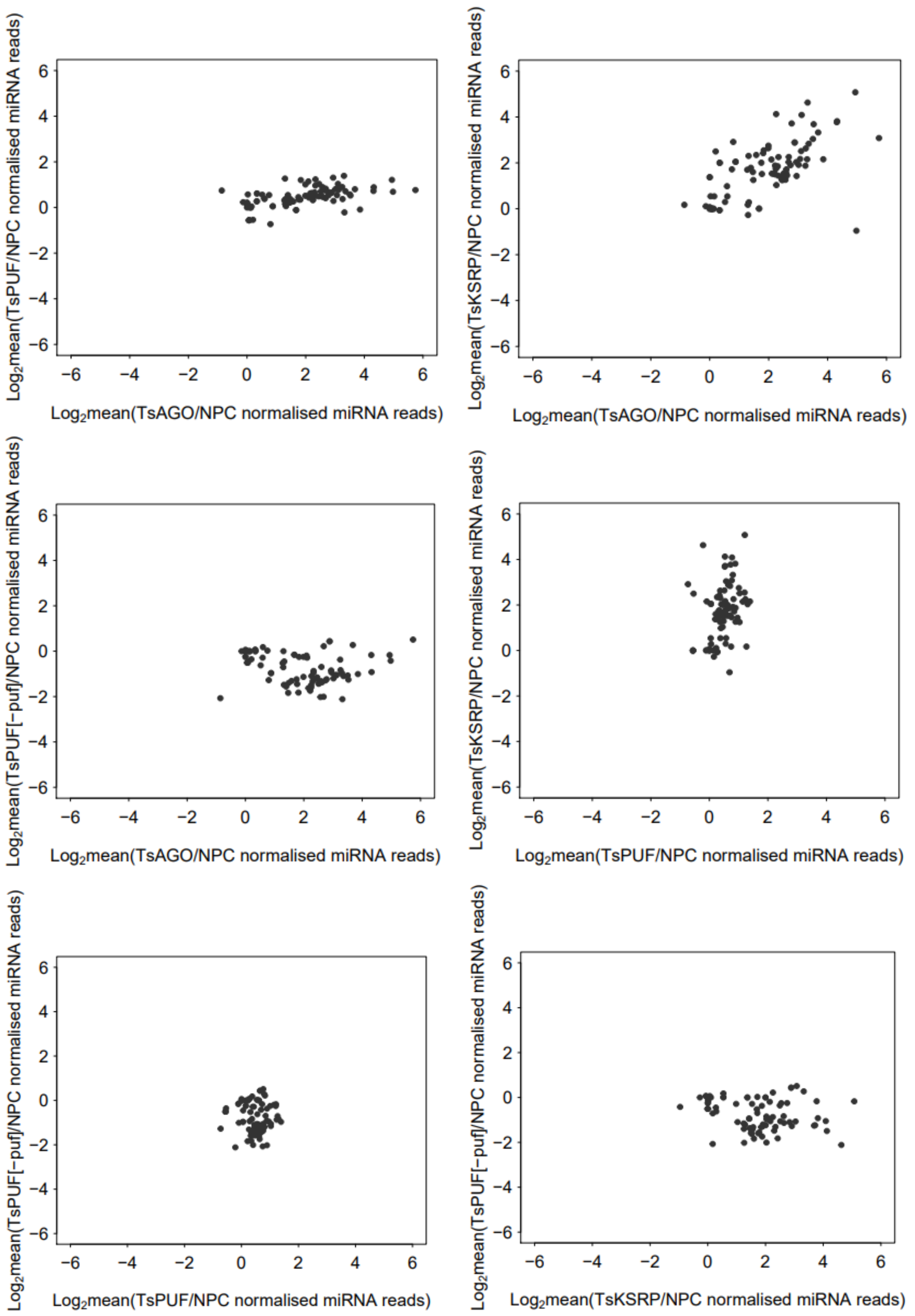


Supplementary Figure 3.

a)

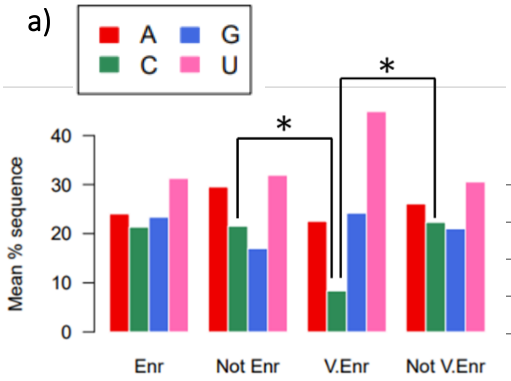


Definitions:

| | |
|---|---|
| Enriched (Enr) | log2(TsKSRP/NPC) > 1 |
| Not enriched (Not Enr) | log2(TsKSRP/NPC) < 1 |
| Very enriched (V.Enr) | log2(TsKSRP/NPC) > 4.5 |
| Not very enriched (Not V.Enr) | log2(TsKSRP/NPC) < 4.5 |

b)

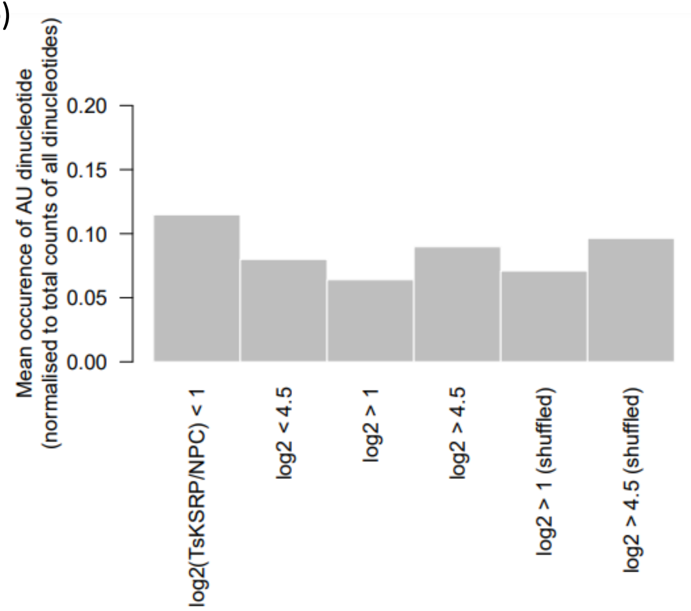


Supplementary Figure 4.

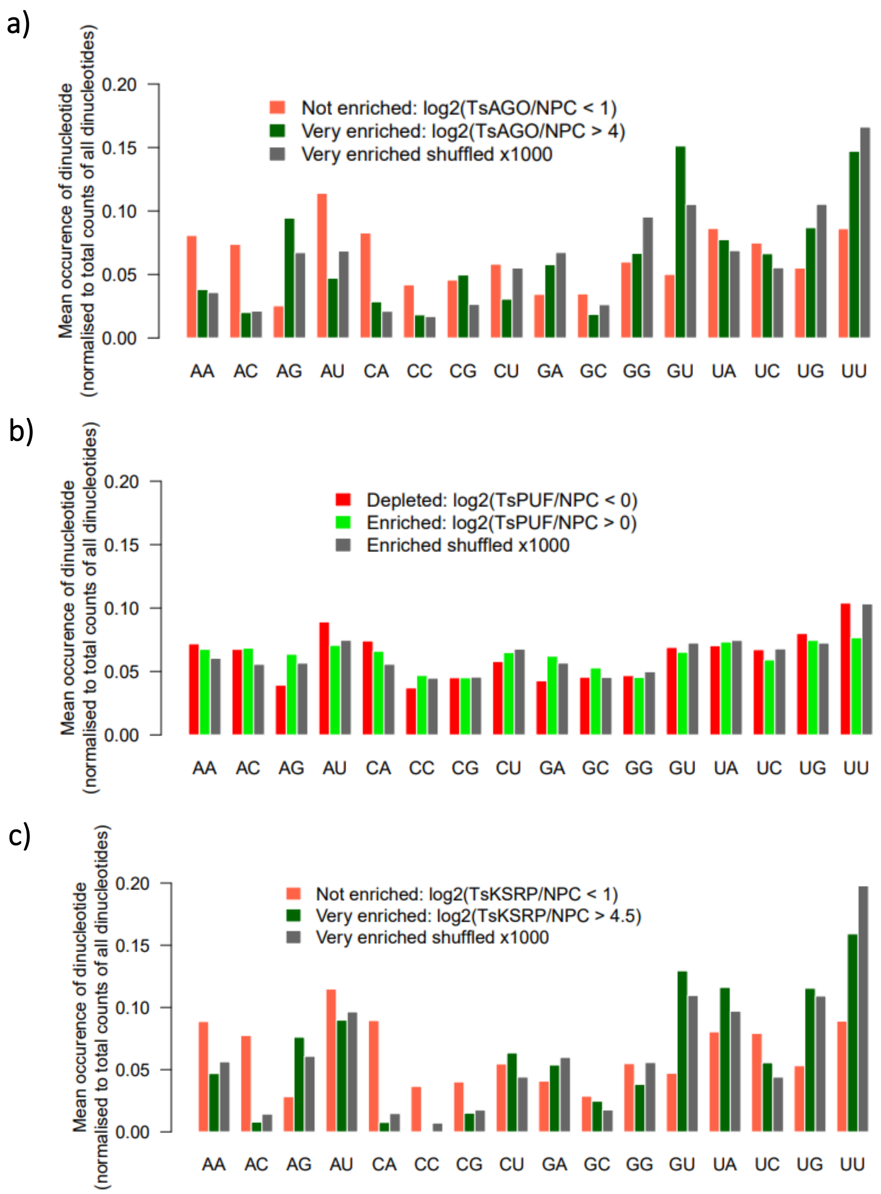


Supplementary Figure 5.

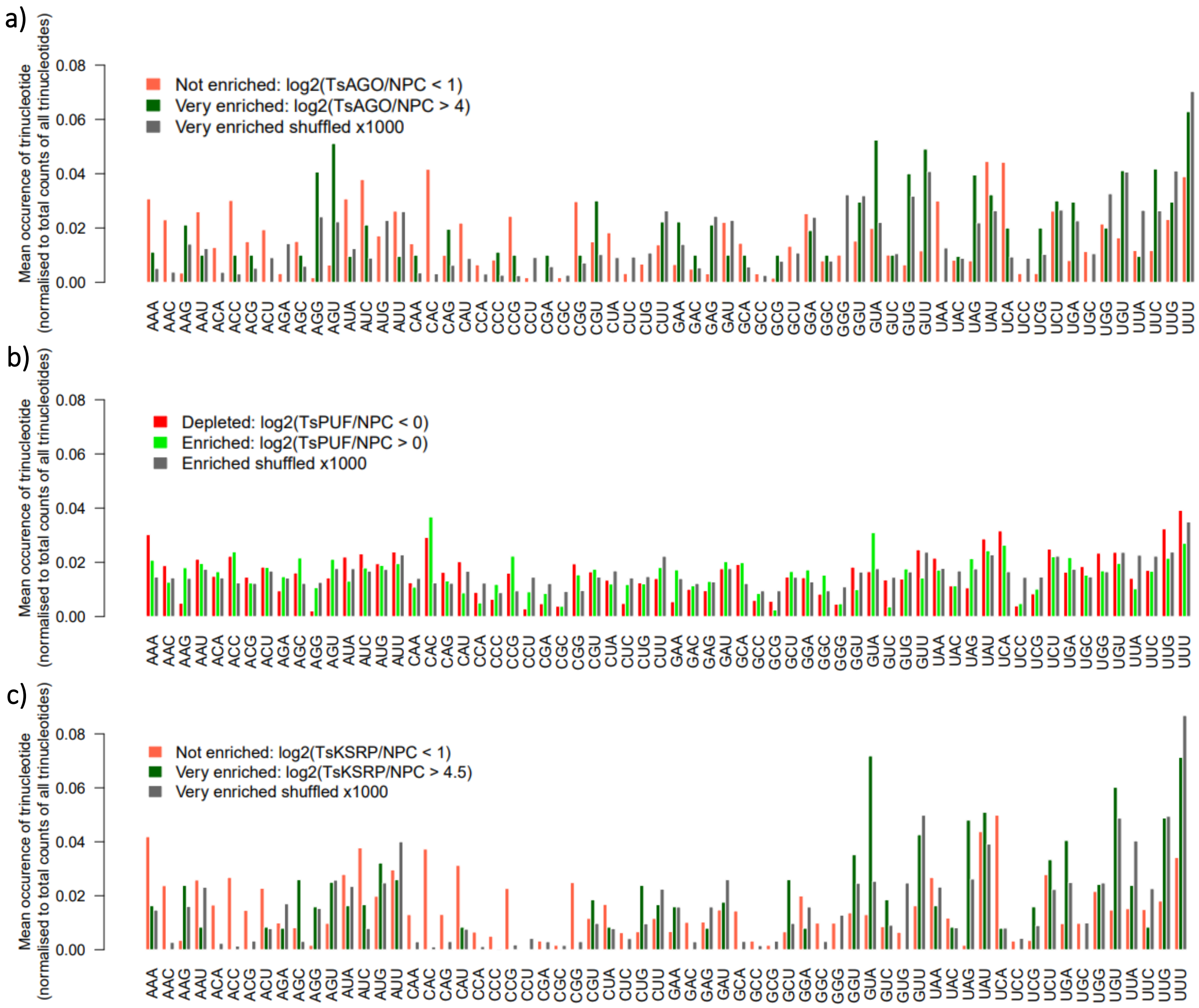


Supplementary Figure 6.

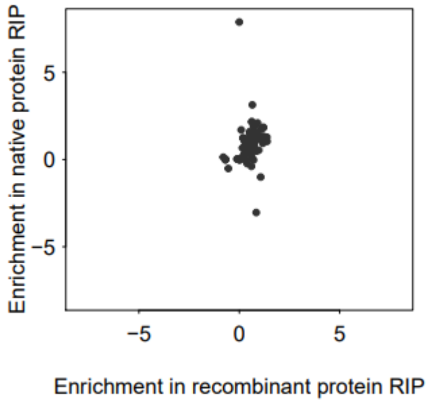


Supplementary Figure 7.

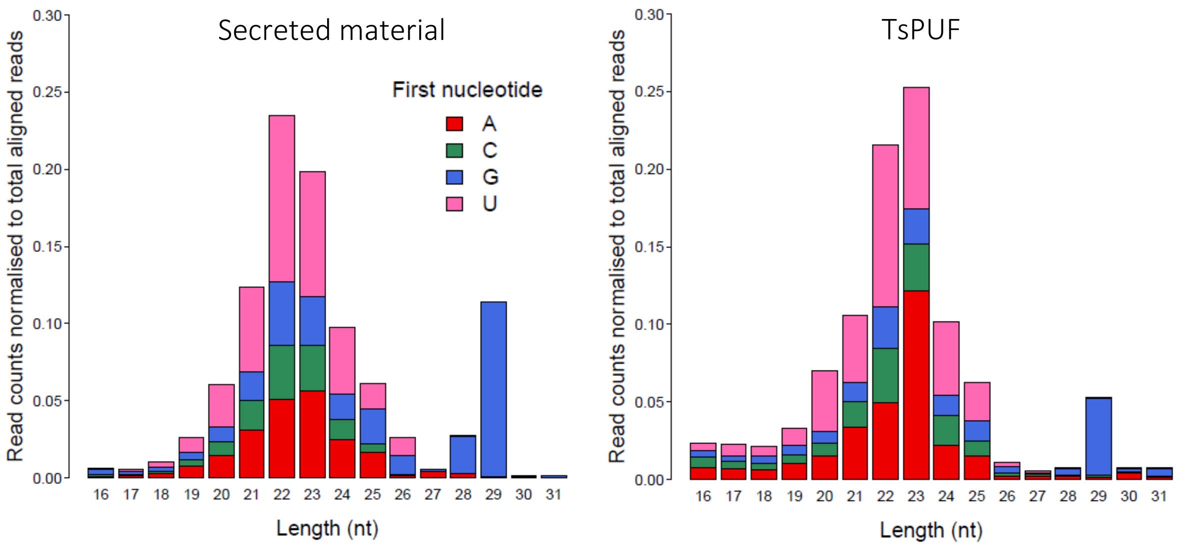


Supplementary Figure 8.
